# Molecular basis of TSC complex GAP activity

**DOI:** 10.64898/2026.07.14.738392

**Authors:** Sonja Titze, Maximilian Rüttermann, Mark Nellist, Daniel Kümmel

**Author notes:** Correspondence (D.K.).

## Abstract

The tuberous sclerosis complex (TSC) protein complex (TSCC) acts as the GTPase activating protein (GAP) for the small GTPase Rheb, to limit mTORC1 (mechanistic target of rapamycin complex 1) activity and cellular growth. We report the structure of the catalytic transition state complex of TSCC and Rheb. Assembly of two TSC2 subunits containing “asparagine-thumb” GAP domains with two accessory TSC1 subunits is required for function in cells. Catalysis requires the “asparagine-thumb” residue of TSC2 and conserved residues in Rheb that bind TSC2 at multiple interaction sites. Surprisingly, only one TSC2 GAP domain is catalytically competent and interacts with Rheb. This is realized by asymmetric binding of TSC1, which enables activating structural changes in one of the TSC2 subunits and locks the second TSC2 copy in an inactive conformation. We identify TSC2 variants that affect conformational coupling within TSCC and binding to Rheb. The structure thus explains the catalytic mechanism of TSCC and reveals an allosteric role of TSC1 in this process.

## Introduction

Small GTPases are molecular switches that regulate cellular processes like signal transduction and vesicular transport, and their dysregulation can cause a variety of diseases ^1^. GTPases either exist in an inactive state when bound to guanosine diphosphate (GDP) or adopt an active conformation in complex with guanosine triphosphate (GTP) to interact with downstream effectors ^2,3^. The cycling between both states is tightly regulated. Guanine nucleotide exchange factors (GEFs) catalyze the exchange of GDP for GTP and thus activate the GTPase. To switch the GTPase off, GTPase activating proteins (GAPs) stimulate GTP hydrolysis and return the GTPase to the GDP-bound state. Because they counteract GTPase signaling, GAPs frequently function as tumor suppressor proteins.

The small GTPase Rheb is an activator of the mTORC1 (mechanistic target of rapamycin complex 1) kinase, the master regulator of cellular growth. Its regulation critically depends on inhibition by the cognate GAP tuberous sclerosis complex (TSC) protein complex (TSCC)^4^. Consequently, this complex is considered a tumor suppressor. Loss of TSCC function is associated with the tumor syndrome tuberous sclerosis complex (TSC)^5^, that affects up to one in 5000 newborns.

TSCC comprises the subunits TSC1, TSC2 (which are conserved in evolution) and a third subunit TBC1D7 in mammalian cells ^6^ (Fig. 1A). TSC2 contains the catalytic ‘asparagine-thumb’ GAP domain homologous to RalGAP ^7^ and RapGAP ^8,9^ and assembles into a tail-to-tail homodimer via a dimerization domain (DD). TSC1 also forms a dimer, consisting of globular N-terminal domains (NTD) that support TSCC function by contributing to lysosomal localization of the complex ^10,11^ and a long C-terminal coiled-coil domain that binds the TSC2 homodimer ^12^. TBC1D7 is a TBC (Tre-2/Bub2/Cdc16) RabGAP domain protein but how this is linked to TSC complex function is not clear ^13,14^. Homozygote or compound heterozygote pathogenic variants in *TBC1D7* cause macrocephaly ^15,16^ and knock-out mice show a partial brain development phenotype (Schrötter et al., 2022), suggesting a specialized role in TSCC regulation.

**Figure 1:**
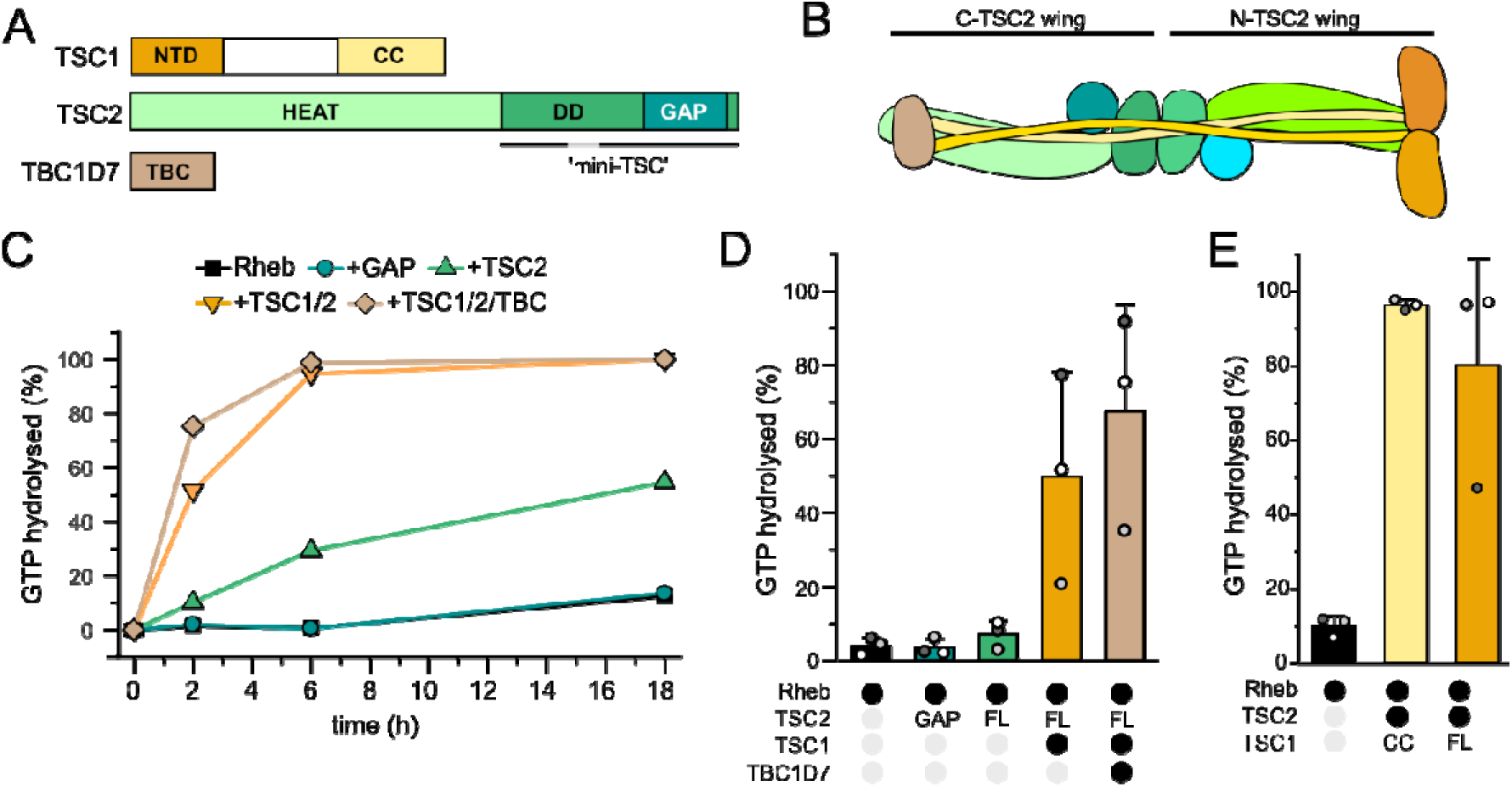
Requirements for TSCC GAP activity. (**A**) Domain architecture of TSC complex subunits. NTD: N-terminal domain; CC: coiled-coil; HEAT: HEAT-repeat domain; DD: dimerization domain; GAP: GTPase activating protein domain; TBC: Tre-2/Bub2/Cdc16 domain. (**B**) Schematic representation of the two-winged TSCC architecture (**C**) Time course of Rheb GTPase activity stimulation by different human TSCC constructs. (**D**) Rheb GTPase activity stimulation by different human TSCC constructs after 2 h. Data from n=3 repeats independent repeats are shown as means ± SD. (**E**) Stimulation of *Cg*Rheb GTP hydrolysis by *Ct*TSC2 with *Ct*TSC1 full-length (FL) or *Ct*TSC1 CC. Data from n=3 independent repeats are shown as means ± SD.

Cryo-EM studies have provided insight into the architecture of TSCC ^11,12,18^ (Fig. 1B). Two copies of TSC2 interact asymmetrically with TSC1, forming a two-winged assembly. However, no structure with the bound substrate Rheb has been reported. Models for the catalytic mechanism of TSC2 rely on mutational analysis and modeling ^19–21^. However, GAP assays showed that full TSC complex has higher activity than the isolated GAP domain ^22^. To understand the molecular basis, we determined the cryo-EM structure of a fully active fungal core TSCC bound to Rheb in the GDP BeF_x_-loaded transition state. The structure provides detailed insight into the catalytic mechanism, uncovers an allosteric contribution of TSC1 to Rheb inactivation, and reveals structural and functional asymmetry of the complex that we validate by mutational studies and the analysis of variants identified in individuals suspected of TSC.

## Results

### TSC1 supports TSCC GAP activity *in vitro*

The isolated GAP domain of TSC2 is in principle capable of stimulating Rheb GTP hydrolysis ^9,19,20^. However, full TSCC or a ‘mini-TSC’ fragment of TSC2 (Fig. 1A) showed higher activity than the TSC2-GAP domain alone *in vitro* ^12,22^, likely arising from interactions of the TSC2-DD domain with Rheb. Furthermore, loss of TSC1 or TBC1D7 in cells resulted in higher Rheb-GTP levels and increased mTORC1 activity ^6,10^. This has been attributed to a requirement of the different domains and subunits for proper TSCC regulation or a role in stabilizing the complex.

We compared the activity of full TSCC to different domains and subcomplexes with a multiple turnover GTP hydrolysis assay (Fig. 1C, 1D, Supplementary Fig. 1). The intrinsic GTP hydrolysis of Rheb was not robustly stimulated by substoichiometric amounts of the TSC2 GAP domain. In contrast, full-length TSC2 clearly accelerated GTP hydrolysis, and was strongly activated by addition of TSC1, leading to complete consumption of GTP in the assay within 6 hours (Fig. 1C). TBC1D7 further increased GAP activity, demonstrating that all three complex subunits are required to reach full catalytic activity of TSCC (Fig. 1D). The greatest relative activity gain was observed upon inclusion of TSC1 in the assay (Fig. 1C, 1D), indicating that in addition to its regulatory function, TSC1 directly contributes to catalytic activity.

Structural studies of human TSCC showed that the TSC1 coiled-coil (CC) domain makes contacts with the TSC2 GAP domain ^11,12^, suggesting that the TSC1-CC domains may be responsible for supporting catalytic activity. Using the TSC proteins form *Chaetomium thermophilium* (*Ct*) and Rheb from *Chaetomium globosum* (*Cg*), we observed that GAP activity of CtTSC2 bound to the CtTSC1-CC was comparable to the CtTSC2-CtTSC1-FL complex (Fig. 1E), demonstrating that the contribution of TSC1 to catalysis is realized by its CC domain.

### Cryo-EM structure of the Rheb bound to TSCC

To understand the combined contributions of TSC1 and TSC2 to GAP activity on a molecular level, we determined the structure of the fully active *Chaetomium thermophilum* core TSCC (cTSCC) bound to Rheb in a GTP hydrolysis transition state. The used expression constructs encoding the *Ct*TSC1 CC domain (residues 550-790) and the HEAT repeat domain (HEAT), dimerization domain (DD) and asparagine-thumb GAP domain (GAP) of *Ct*TSC2 (residues 1-1520) (Supplementary Fig. 2A). Reconstitution of a TSCC-Rheb complex failed, likely due to instability of the transient GAP-GTPase complex. We thus fused *Cg*Rheb to the C-terminus of TSC2 (Supplementary Fig. 2A), which is an established approach to study the catalytic mechanism of GAP-GTPase complexes ^23,24^. The fusion construct was stable and active in GTP hydrolysis assays (Supplementary Fig. 2B, 2C), demonstrating that the linker did not interfere with GAP function. To capture the catalytic transition state of GTP hydrolysis for structural analysis and to stabilize cTSCC-Rheb interactions, the purified complex was incubated with GDP and BeF_x_ prior to EM grid preparation.

For cryo-EM data collection we recorded untilted and 40° tilted data to compensate for preferential orientation of the particles (Supplementary Table 1). During single particle analysis, we identified two species in the samples (Supplementary Fig. 3, 4) with about half of the particles showing additional density at the center that likely represents bound Rheb (Supplementary Fig. 3). Separate processing yielded 2D classes of apo cTSCC and Rheb-bound cTSCC. 3D reconstruction was complicated by the elongated shape and high intrinsic flexibility of the particle, resulting in poor resolution at the tips of the structure (Supplementary Fig. 4). Applying a mask centered at the TSC2 dimerization interface and implementation of EMready2 ^25^ resulted in maps at 3.06 Å and 3.01 Å resolution (gold standard FSC_0.143_ ^26^) of apo-cTSCC and Rheb-cTSCC, respectively (Fig. 2A-D). To improve resolution of the additional density from bound Rheb, we also applied a focused mask to this region and performed local refinement.

**Figure 2:**
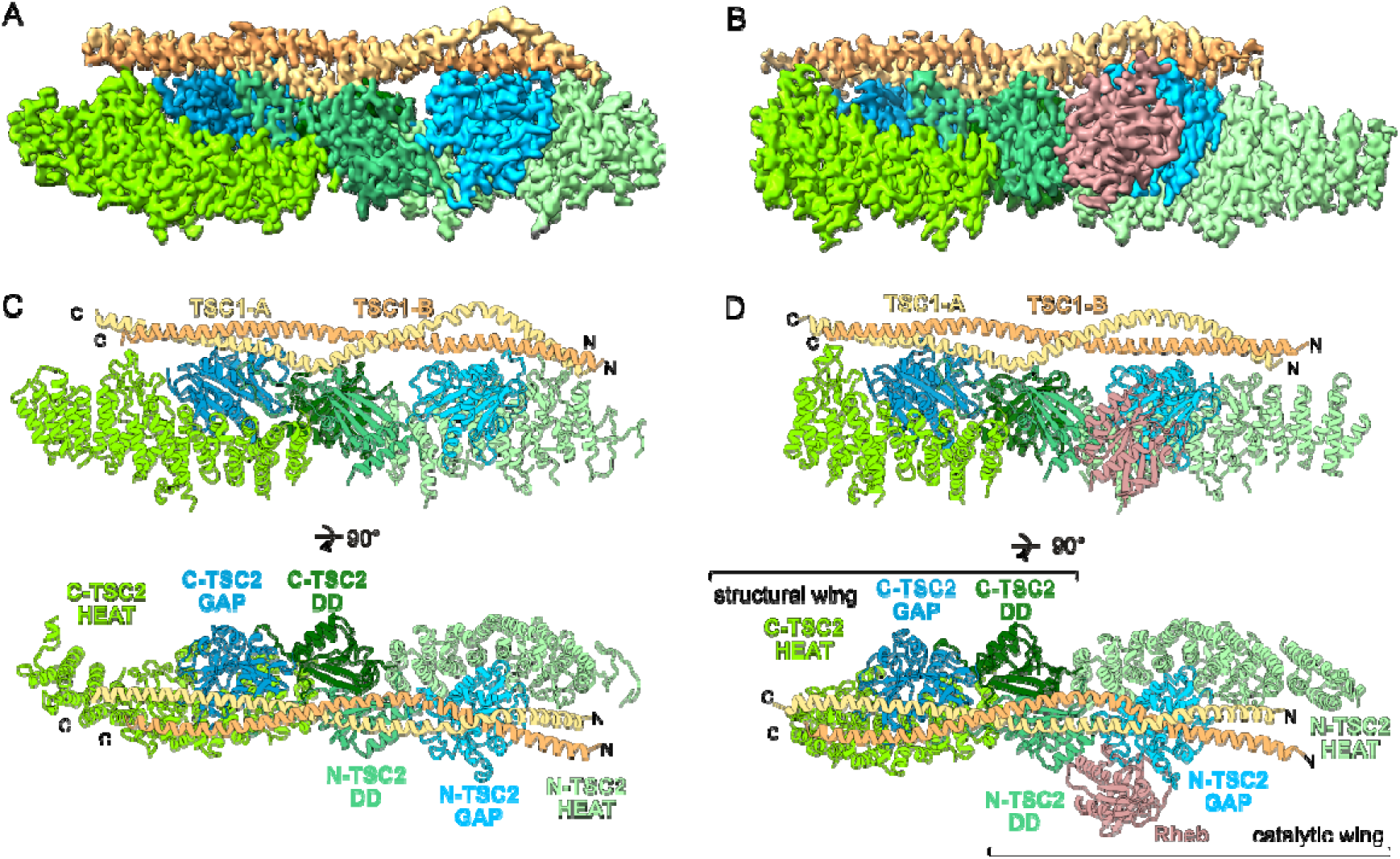
Structure of apo cTSCC and Rheb-boud cTSCC. (**A**) Coulomb potential maps of the apo cTSCC reconstruction and (**B**) Rheb-bound cTSCC reconstruction. (**C**) Orthogonal views of the models of apo cTSCC and (**D**) Rheb-bound cTSCC.

This allowed us to unambiguously fit and rebuild models of *Ct*TSCC in both maps. *Ct*TSC2-GAP, -DD, and the C-terminal portion of *Ct*TSC2-HEAT were well resolved. The *Ct*TSC1 CC was less well defined but backbone tracing was possible. By combining the model from cryo-EM with the X-ray structure of the *Ct*TSC2 N-terminal domain ^27^, we obtained a model of the entire *Ct*TSCC that fits well with the 2D class averages and shows a characteristic elongated shape of ∼30 nm length (Supplementary Fig. 4G), highly similar to human TSCC. The subunit architectures are also conserved between fungal and human TSC proteins. The TSC2 HEAT domain is followed by a dimerization domain that mediates tail-to-tail interaction of two TSC2 subunits (Supplementary Fig. 5A). The C-terminal GAP domain comprises an eight-stranded β-sheet and four helices, with α3 referred to as catalytic helix as it contains the essential asparagine-thumb residue (Supplementary Fig. 5B). The GAP domain folds back onto TSC2-DD and TSC2-HEAT, resulting in a tripartite arrangement (Fig. 1B, Fig. 2).

### Conformational differences between the two TSC2 subunits

The TSC1-CC dimer runs alongside the HEAT and dimerization domains of both TSC2 subunits and induces asymmetry in the complex (Fig. 2, 3A, 3B). In the following, we refer to the TSC2 subunits that are associated with the N-terminal and C-terminal portions of the TSC1 coiled coil as N-TSC2 and C-TSC2, respectively. The asymmetry is not dependent on Rheb binding and clearly resolved in the structures of both apo cTSCC (Fig. 3A) and Rheb-bound cTSCC (Fig. 3B). (i) One notable difference between the two copies of TSC2 is a loop region between β4 and αA of the dimerization domains (D-loop) that is only resolved in N-TSC2 but disordered in C-TSC2 (Fig. 3C). The N-TSC2 D-loop reaches over to the GAP domain of C-TSC2 and the C-terminal portion of TSC1-CC. Thus, the interaction of TSC1 and C-TSC2 is stabilized by the D-loop. (ii) The linker sequence connecting the HEAT and dimerization domains (HD-linker) of TSC2 occupies different binding sites in the two wings. In C-TSC2, a short helical segment in the linker binds to a groove between the GAP and dimerization domain. In N-TSC2, this segment interacts with TSC1 instead and is displaced by ∼27 Å (Fig 3D). (iii) Furthermore, the loops connecting strands β2 and β3 in the GAP domains of C-TSC2 and N-TSC2, referred to as the β2-β3 loop hereafter, adopt slightly different conformations (Fig. 3E).

**Figure 3:**
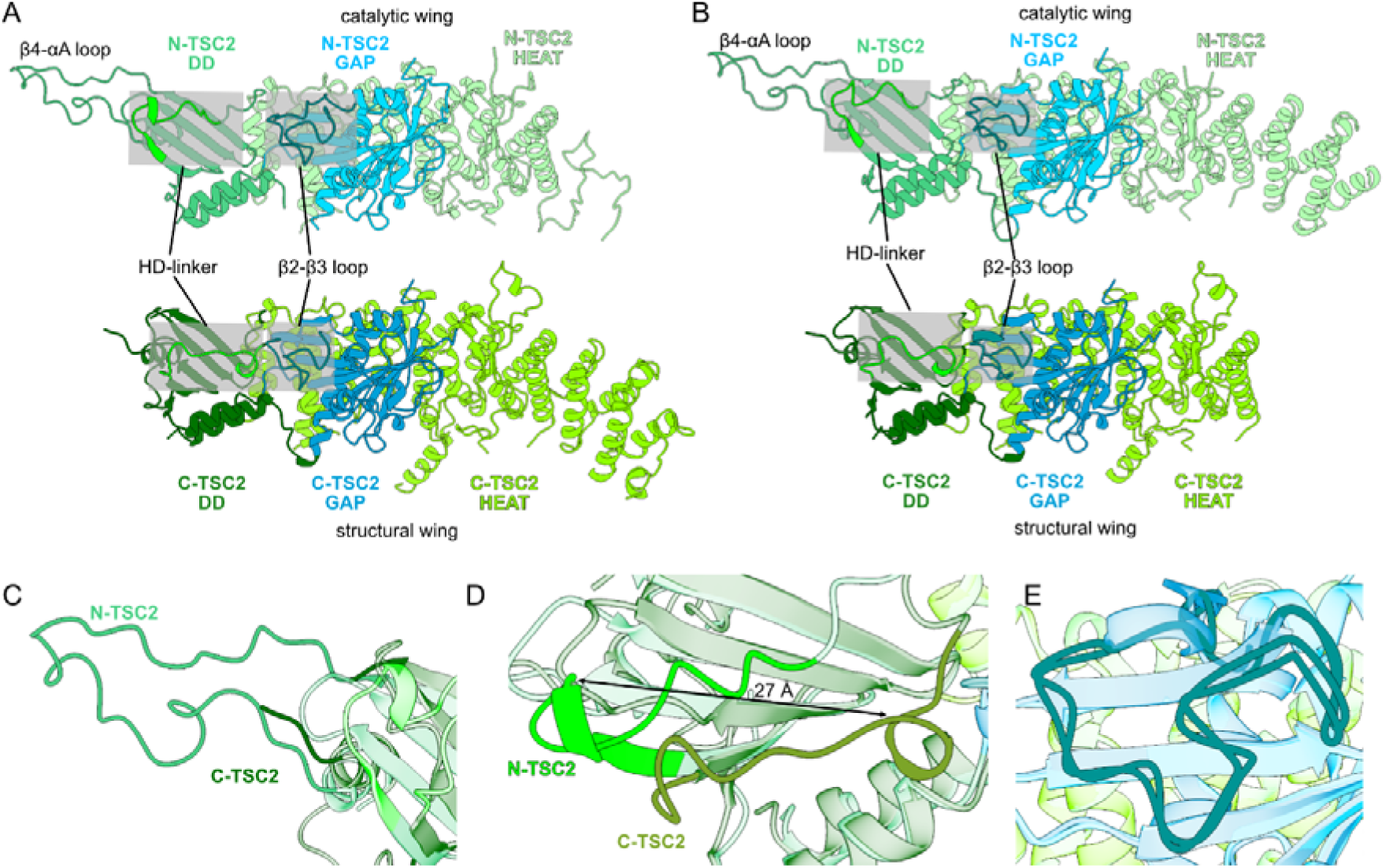
Asymmetry of TSC2 in cTSCC. (**A**) Comparison of N-TSC2 (top) and C-TSC2 (bottom) in apo cTSCC and (**B**) Rheb-bound cTSCC. (**C**) Close-up of an overlay of the β4-αA D-loop region, (**D**) the HD-linkers, and (**E**) the β2-β3 GAP-loops of N-TSC2 versus C-TSC2.

### Concomitant stabilization of inactive and active GAP domain conformations by TSC1

Surprisingly, additional density representing bound Rheb is only associated with N-TSC2 (Fig. 2C, 2D) and we did not identify particles with two Rheb molecules bound. Using symmetry expansion, we confirmed that this observation reflects a feature of the particles and not a data processing artefact (Supplementary Fig. 6, see Methods section for details). Rheb could be modelled with a Mg^2+^ ion, a GDP molecule, and BeF_3_ at the γ-phosphate position occupying the nucleotide binding pocket (Supplementary Fig. 4H). The structure thus represents the catalytic active GTP hydrolysis transition state of the cTSCC-Rheb complex.

We wondered why the two GAP domains in TSCC are not equally competent in engaging Rheb to promote GTP hydrolysis and reasoned that the asymmetry of N-TSC2 and C-TSC2 introduced by TSC1-CC binding may provide an explanation. Compared to the apo structure, N-TSC2 undergoes structural rearrangements at three regions to enable Rheb binding (Fig. 4A): (i) The relative orientation of the DD and GAP domain changes, causing two N- and C-terminal helical extensions (αN and αC, Supplementary Fig. 5A) that are part of the DD to move 4-4.5 Å closer to Rheb and interact with the GTPase (Fig. 4B). (ii) The β2-β3 loop of the GAP domain undergoes conformational changes that affect side chain positioning (Fig. 4C). (iii) The TSC1 CC domains change their positioning on N-TSC2 (Fig. 4D), moving ∼7.5 Å further away from the GAP domain.

**Figure 4:**
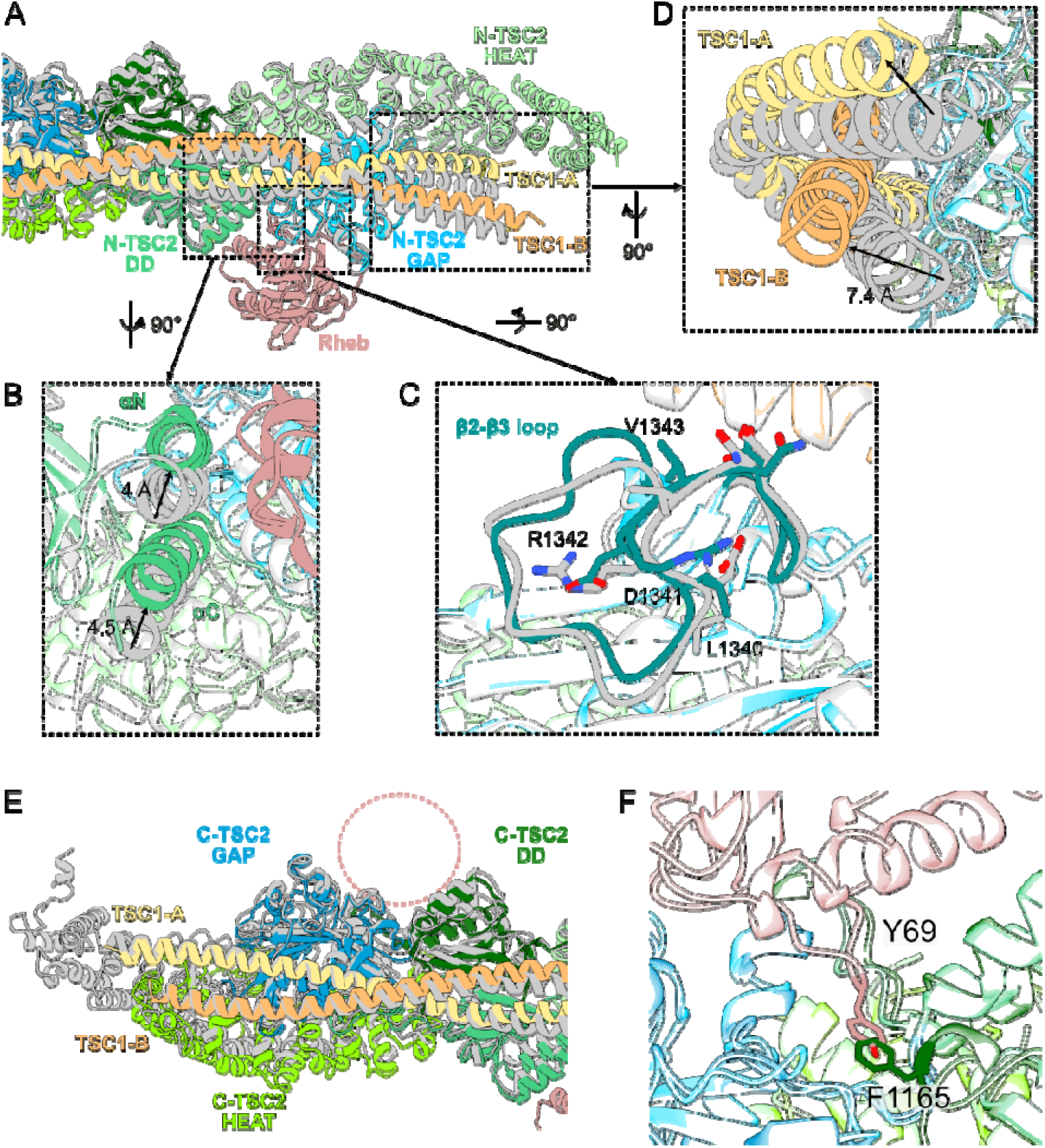
Conformational changes of cTSCC accompanying Rheb binding. (**A**) Superposition of the N-TSC2 catalytic wing of apo cTSCC (grey) and Rheb-bound cTSCC (colored as in Figure 2). (**B**) Close-up of the αN/αC helices of the TSC2 DD of apo (grey) and Rheb-bound cTSCC (green). (**C**) Close-up of the TSC2 GAP domain of apo (grey) and Rheb-bound cTSCC (cyan). (**D**) Close-up of the TSC1-CC domains associated with apo (grey) and Rheb-bound cTSCC (yellow/orange). (**E**) Superposition of the C-TSC2 structural wing of apo cTSCC (grey) and Rheb-bound cTSCC (colored as in Figure 2). The proposed Rheb binding site on C-TSC2 is indicated by a pink sphere. (**F**) Modeling of Rheb interaction with the C-TSC2 GAP domain based on its position at N-TSC2. The conformation of the C-TSC2 HD-linker prevents binding of Rheb at the structural wing.

Thus, Rheb binding requires concerted structural rearrangements of the TSC2 and TSC1 subunits in one half of the complex that represents the “catalytic wing”. These conformational adjustments are also observable in the symmetry expansion analysis (Supplementary Fig. 6, Supplementary Movie 1). Importantly, the participation of TSC1 in formation of the Rheb catalytic site by allosteric coupling provides a molecular explanation of why TSC1 is required for full GAP activity of the TSCC.

Comparing the models of C-TSC2 and the associated part of TSC1 in the apo and Rheb-bound cTSCC structures we observe virtually identical conformations (Fig. 4E). This wing of the complex is stabilized by the ordered D-loop of N-TSC2 that reaches over onto the C-TSC2 GAP domain (Fig. 3C). In this conformation C-TSC2 is apparently inactive and cannot bind Rheb, which renders this half of the complex a merely “structural wing”. Docking Rheb to the GAP domain of C-TSC2 by superposition of the catalytic wing revealed a clash between F1165 in the HD-linker of C-TSC2 with Y69 of Rheb (Fig 4F). This shows that binding of Rheb to the structural wing is prevented by the specific conformation of C-TSC2.

### Rheb engagement via multiple sites is required for TSCC activity

Rheb binding to TSC2 occurs via (i) the catalytic α3 helix, (ii) the β2-β3 loop of the GAP domain, and (iii) the αN/αC helical extensions that are part of the DD (Fig. 5A). The α3 helix interacts with the Rheb switch I region and inserts its asparagine thumb residue (N1388) into the nucleotide binding pocket. The active site of the Rheb-TSCC structure is clearly resolved in the reconstruction (Supplementary Fig. 4H, 4I). GDP•BeF_3_ is bound in the nucleotide binding pocket, mimicking the GTP hydrolysis transition state, and TSC2 N1388 is oriented towards the nucleotide at a distance of 3.9 Å to the BeF_3_ (Fig. 5B). Likewise, the conserved switch I residue Y37 is positioned in the active site at hydrogen bonding distance to BeF_3_. This residue (Y35 in human RHEB) is critical for GAP function ^21^ and, interestingly, pathogenic *RHEB* variants causing amino acid substitutions at codon 35 have been found in individuals with TSC-like clinical manifestations ^28^. This residue is also conserved in the GTPase Rap where it was reported to stabilize formation of the GTP hydrolysis state in the complex with RapGAP ^9^. Overall, the arrangement of the Rheb-TSC2 active site is similar to the Rap-RapGAP complex (Fig. 5C). The structure supports a model for the catalytic mechanism of TSCC that involves the asparagine-thumb residue as the sole catalytic residue in TSC2 and requires support from a tyrosine in Rheb that stabilizes formation of the transition state ^19^.

**Figure 5:**
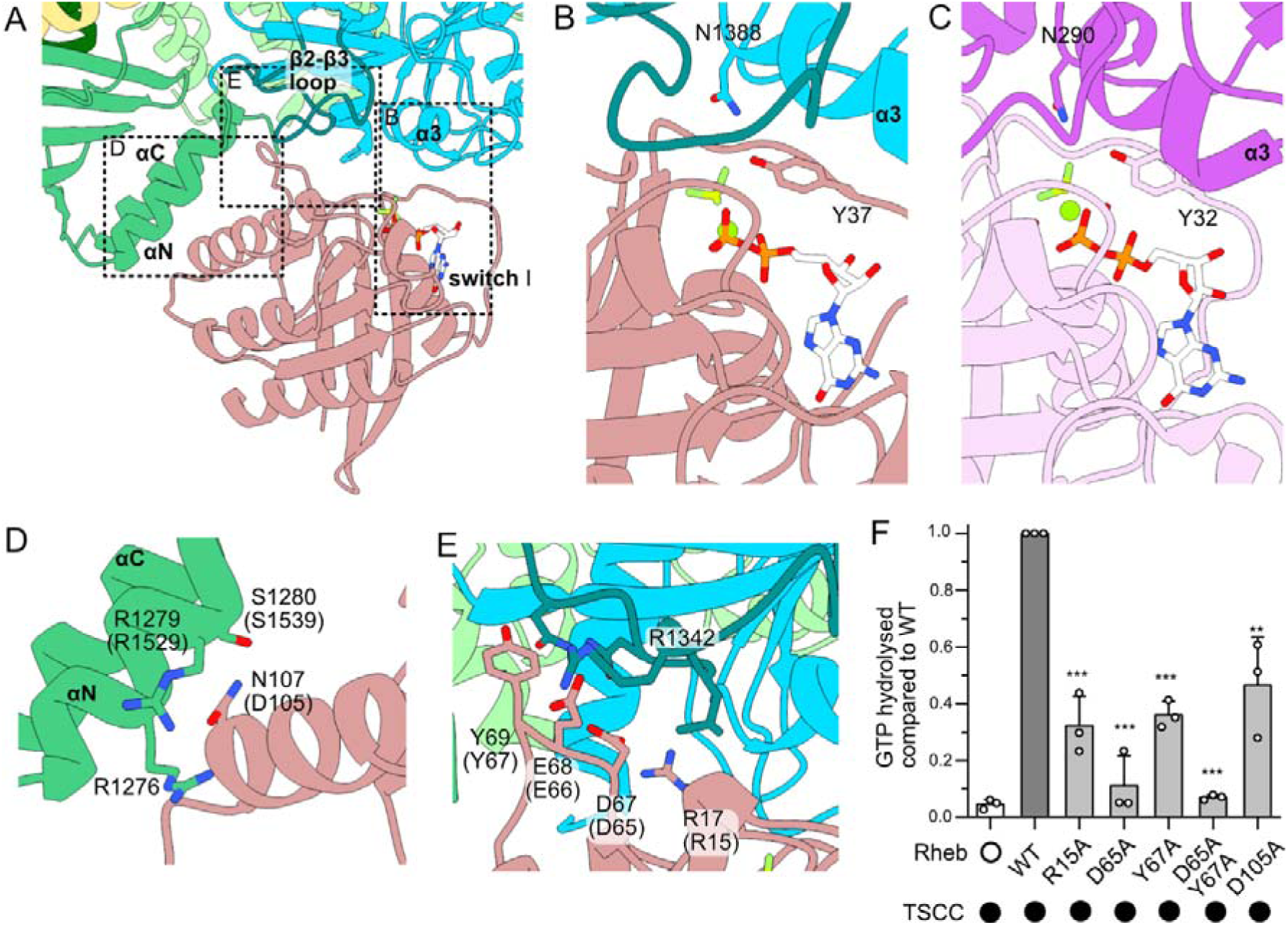
Rheb interactions with TSC2. (**A**) Overview of key interaction interfaces between Rheb and TSC2. (**B**) Close-up of the Rheb-TSC2 active site. (**C**) Close-up of the Rap-RapGAP active site (PBD ID 3BRW, ^9^). (**D**) Interaction site of Rheb N107 with TSC2 αN/αC helices. (**E**) Interface between the TCS2 β2-β3 loop and Rheb. Residues in brackets denote the equivalent positions in human homologs. (**F**) GAP assays with human TSCC and Rheb mutants. Data from n=3 repeats independent repeats are shown as means ± SD. Statistical analysis with pairwise t-test, ** p < 0.001, *** p < 0.0001.

In addition to the active site, secondary binding sites for Rheb are created by the structural changes in N-TSC2 (Fig. 3C, 3D). The two helices of the TSC2-DD, leading into (αN) and out of (αC) the GAP domain provide an additional interaction site for Rheb (Fig. 5D) that was proposed before based on the homology to RapGAP ^9,12^. The interface is formed around N107 of Rheb, which is cradled between αN and αC. This interaction is only possible after the conformational rearrangement of N-TSC2 (Fig. 3D). The rearrangement of the β2-β3 loop creates a binding pockets for Rheb R17 and D67, which are engaged in an intramolecular salt bridge, and Rheb E68 (salt bridge with TSC2 R1342) (Fig. 5E). Additionally, Y69 interacts with the β2-β3 loop and makes contacts to TSC2-DD (Fig. 5E). This binding site for Y69 is blocked in the inactive TSC2 copy of the structural wing (Fig. 4G), suggesting that the interaction may be essential for productive engagement of Rheb by TSC2.

To test the relevance of the secondary Rheb interactions with TSC2 for function, we mutated key residues in the interfaces and tested the effect on GAP activity (Fig. 5A, 5F). The residues of interest are conserved between *Cg*Rheb and the human isoforms RHEB and RHEBL (Supplementary Fig. 7A). Functional assays were therefore performed with the human proteins. TSCC-stimulated GTP hydrolysis was strongly reduced by D105A substitution in the center of the Rheb interface with αN/αC of the TSC2-DD (Fig. 5D, 5F). The R15A and D65A substitutions were detrimental to GAP activity of TSCC (Fig. 5E, 5F), recapitulating the effect on activity observed with the isolated GAP domain ^20,21^. This was also observed for the Y67A substitution, which affects the interactions with both the β2-β3 loop of the GAP domain and αN of the DD. This finding corroborates the conclusion that blockage of the Rheb tyrosine binding site of C-TSC2 is the key structural feature that renders the structural wing catalytically inactive. Collectively, the data suggests that full TSCC complex activity requires multiple TSC2-Rheb interaction sites that emerge in the catalytical wing and are prevented in the structural wing through TSC1-supported structural rearrangements.

### Allosteric coupling in TSCC conveys GAP activity

The analysis of the cryo-EM reconstruction suggests that TSCC GAP activity requires concerted conformational changes of TSC1-CC, TSC2-GAP and TSC2-DD. The β2-β3 loop plays a central role in relaying the structural rearrangement as it contacts the TSC1-CC domains, the catalytic helix α3, and the αN/αC helices of the TSC2-DD. We reasoned that perturbations in the key elements of TSC2 that convey allosteric communication should impair TSCC function. To test this hypothesis, we investigated 10 human *TSC2* variants that have been reported in individuals suspected of TSC and map to the αN/αC helices and β2-β3 loop (Fig. 6A, Supplementary Table S2) ^29,30^. The variants were tested for their ability to restore mTORC1 inactivation in *TSC1^-/-^*/*TSC2^-/-^*double knockout cells, compared to wild-type (WT) TSC2 and the pathogenic TSC2 R611Q variant that shows reduced TSC1 binding (Fig. 6B, 6C). This assay is well established for assessing the pathogenicity of *TSC2* variants and to obtain mechanistic insight into TSCC function ^31^. The variants tested expressed at similar levels to wild-type, suggesting that overall structure and stability was not affected (Supplementary Fig. 8A). Furthermore, levels of co-expressed TSC1 were comparable, which indicates intact complex assembly (Supplementary Fig. 8B). Only two variants, R1529W and Q1532R reduced mTORC1 activity to a similar level as WT TSC2. These residues map to the interaction interface of the TSC2-DD helix αN with Rheb. Interestingly, the mutations L1534H and D1535A that contribute to the stability of the αN/αC region of the DD inactivated TSCC, suggesting that structural integrity is more important than surface properties. (Fig. 6D). All variants in the β2-β3 loop negatively affected TSC2 activity. Mutation of L1594, G1595, and G1596 to bulky polar residues is expected to disrupt the folding of the loop (Fig. 6E). D1603 connects the β2-β3 loop with the catalytic α3 helix via a salt bridge to R1639 (Fig. 6F). Comparing the two variants tested for G1596 (to alanine or aspartate) and D1603 (to glutamate or valine), respectively, the effects on mTORC1 inhibition *in vitro* correlate with the degrees of chemical difference resulting from the exchange.

**Figure 6:**
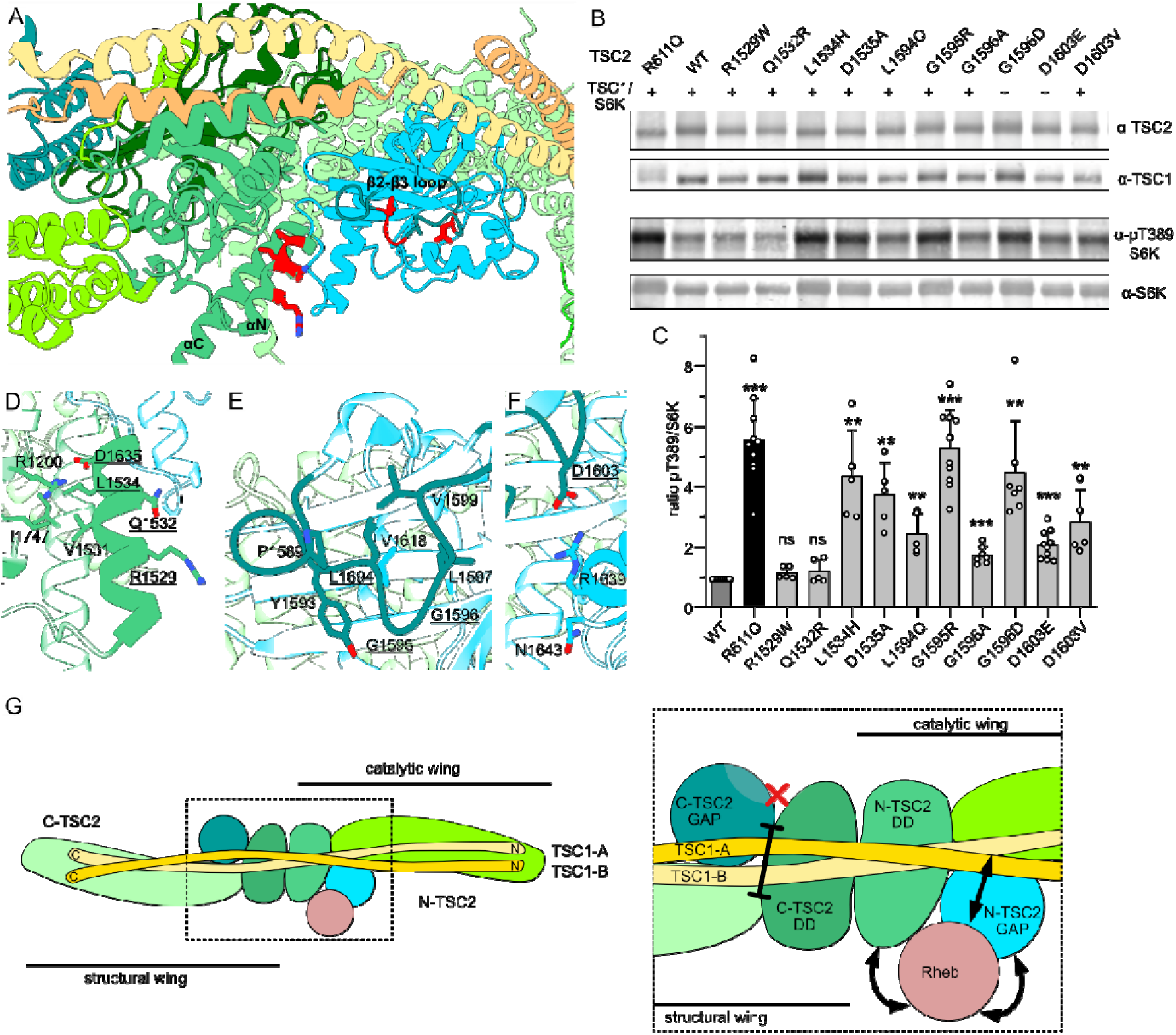
Characterization of TSC2 variants in the dynamic Rheb interface. (**A**) TSC2 variants mapped in the cryo-EM structure of TSCC (PDB ID 9CE3 ^11^). (**B**) Functional assessment of variants with a cellular mTORC1 activity assay. (**C**) Quantification of pS6K/S6K levels from B. Data from n=5-10 repeats are shown as means ± SD. Statistical analysis with pairwise t-test, ** p < 0.01, *** p < 0.001. (**D**) Interactions of tested TSC2 variants in DD αN helix. (**E**) Mutations in the TSC2 GAP domain β2-β3 loop are expected to interfere with folding. (**F**) D1603 links the β2-β3 loop to the catalytic helix α3 via a salt bridge. (**G**) Schematic summarizing the inhibitory (structural wing) and activating (catalytic wing) allosteric control of TSC2 by TSC1.

Thus, structural integrity of the β2-β3 loop and the αN/αC helices is essential for TSCC function, while activity is less vulnerable to interface variants. This demonstrates that the association of TSC2 with Rheb is more sensitive to structural perturbations than to changes in surface properties.

Taken together, the functional data are consistent with a model that allosteric remodeling of TSC1 and TSC2 for Rheb binding is a prerequisite for TSCC GAP activity. Defects in mTORC1 regulation caused by variants that disrupt the conformational landscape of TSC2 were identified in individuals suspected of TSC. This supports the physiological relevance of this mechanism.

## Discussion

The structural and functional analysis of TSCC presented here provides in depth insight into the Rheb GAP mechanism with important implications. First, our results confirm a proposed catalytic mechanism ^19^ that requires the asparagine-thumb residue provided by TSC2 and a conserved tyrosine in Rheb switch I. Rheb residues D65 and R15, also previously proposed as catalytic residues ^21^, are not part of the active site but bind to TSC2 at secondary sites. Unexpectedly, coordinated structural rearrangements of both TSC1 and TSC2 are required for engagement of Rheb in a GAP-competent state at one wing of TSCC.

In contrast, the second TSCC wing is rigidified in a conformation that is incompatible with Rheb binding. It was unexpected that only one GAP domain of the two TSC2 subunits in the TSC complex is enzymatically competent. Although symmetry in structures of TSCC was previously observed at a global scale ^11,12^, the limited resolution and map quality at the GAP domain interfaces did not allow to resolve the conformational differences at amino acid side chain level. Our structure clearly demonstrates that the different architectures of the two TSC2 wings result in large conformational variations in the dimerization and GAP domains that have striking functional consequences.

The structure indicates that the asymmetry is imposed through binding of the TSC1 CC dimer along the TSC2 homodimer. The TSC1-induced asymmetry results in higher flexibility of the GAP domain β2-β3 loop of N-TSC2. In contrast, folding of the N-TSC2 D-loop onto TSC1 and C-TSC2 likely reduces the flexibility in this part of the complex. This is reinforced by divergent TSC2 HD-linker conformations that prevent Rheb binding specifically in the structural wing. Thus, TSCC is designed to contain a more rigid GAP-inactive structural wing and a GAP-active catalytic wing with higher structural plasticity (Fig. 6G).

Fittingly, the lipid binding sites of TSC1 and WIPI are associated with the catalytic wing and the model of membrane recruitment as proposed by Bayly-Jones *et al*. ^11^ are compatible with our findings: the GAP domain of the catalytic wing is predicted to face the membrane and could reach Rheb. By contrast, the short hyper-variable domain of Rheb would prevent interaction with the structural wing that is oriented towards the opposite side. Rending the structural wing catalytically inactive may therefore safeguard TSCC from adopting competing productive and non-productive lysosomal recruitment modes. Although speculative, this could be relevant if a defined orientation on the organelle membrane is required for proper communication between TSCC and other components of the mTORC1 signaling network.

We can only speculate on potential alternative functions of the structural wing at this point, but it is interesting to note that TSCC was also reported to bind inactive GDP-Rheb, RhoB and Rag GTPases ^14,32,33^. The GAP domain of the structural wing could thus represent a binding site for one or more of these GTPases, possibly mediating different recruitment modes of TSCC to the lysosome.

Our findings provide a molecular explanation for the requirement of TSC1 for GAP activity of TSCC and add a new role to the list of TSC1 functions. TSC1 stabilizes TSC2 ^34^, which we also observed during recombinant protein production, where we obtained lower yields for TSC2 when expressed alone than when co-expressed with TSC1 (Supplementary Fig. 1A). Furthermore, TSC1 is required for recruitment of TBC1D7 ^35–37^, which mediates a specialized function of TSCC in neurodevelopment that is not fully understood ^17^. More recently, we showed that TSC1 helps localize the TSCC to lysosomes ^10^. The manifold contributions of TSC1 to stability, localization and activity of TSCC explain why loss of TSC1 causes TSC, while the peripheral subunits TBC1D7 and WIPI3 do not ^6,38^.

The direct contribution of TSC1 to catalytic activity is reminiscent of the mechanism we recently described for the homologous RalGAP complexes ^7,39^. The RalGAPα subunit shares a conserved domain architecture with TSC2, including an asparagine-thumb GAP domain that alone is catalytically inactive. Binding of the RalGAPβ subunit is required for activity and involves the association of a RalGAPβ helix with the RalGAPα GAP domain. This interaction site is equivalent to the interface between TSC1 and the TSC2-GAP domain. Thus, although TSC1 and RalGAPβ are structurally unrelated, they fulfil conserved functions in enabling catalytic activity of their cognate asparagine-thumb domains via similar structural motifs.

A better understanding of TSCC function will be helpful in determining the likely pathogenicity of *TSC2* variants of uncertain clinical significance (VUS). Combining structural and functional characterization, we here identify VUS that impair TSC complex function *in vitro* and therefore could contribute to TSC or TSC-like pathologies. Importantly, the improved insight into the catalytic mechanism will also be instrumental for defining minimal TSC2 and TSC1 constructs that can be used in gene therapy ^40,41^.

## Acknowledgments

We are grateful to Angela Peron (Division of Medical Genetics, Meyer Children’s Hospital IRCCS, University of Florence, Italy) and Luciana Haddad (University of Sao Paulo, Brazil) for communicating unpublished *TSC2* variants. This work was supported by grants to D.K. from the German Research Council (DFG KU2531/3) and the US Department of Defense TSC Research Program (TS190029). Cryo-EM data were collected at the “Cryo-EM SoN” infrastructure of the University of Münster, funded by the DFG (project number 496113311). We are grateful to Alexander Neuhaus for valuable user support. Cryo-EM data processing was carried out on the Palma II HPC cluster (DFG INST 211/667-1) of the University of Münster. We thank Ann-Marie Lawrence-Dörner and Sabine Steidel for excellent technical assistance and all members of the Kümmel Lab for helpful discussions.

## Materials availability

Further information and requests for resources and reagents should be directed to and will be fulfilled by the corresponding author.

## Data availability

Atomic coordinates and EM maps have been deposited into the protein database (apo core TSCC - PDBID: 31AX, EMD-58238; Rheb-bound core TSCC - PDBID: 31AW, EMD-58237).

## Methods

### Cloning and Mutagenesis

*Chaetomium thermophilum TSC1* and *TSC2* (Genscript) and *Chaetomium globosum* Rheb (GenArt,Thermo Scientific) were amplified from codon-optimized synthetic genes using Q5 Polymerase (NEB). Mutations were introduced with the Q5 (NEB) or QuikChange II XL (Agilent) site directed mutagenesis kits. The integrity of all constructs (listed in Supplementary Table 3) was verified by sequencing.

### Protein Production with Bacteria

BL21 *E. coli* cells transformed with Rheb and TSC2-GAP (human and *Ct*) expression plasmids were grown to an OD_600_ of 0.6-0.8 and protein expression was induced with 0.5 mM IPTG. Proteins were produced overnight at 16°C and cultures harvested by centrifugation (3500 rpm, 15 min) were used directly or stored at -70°C. Cell pellets were lysed in PBS (Rheb, 100 mM Na_2_HPO_4_ pH7.4, 137 mM NaCl, 27 mM KCl, 18 mM KH_2_PO_4_) or GAP Affinity buffer (50 mM NaH_2_PO_4_ pH 8, 500 mM NaCl, 5% Glycerol) supplemented with 1 mg ml^-1^ lysozyme, 0.025 mg ml^-1^ DNAse, 1× protease inhibitor cocktail (PIC, HP, Serva Electrophoresis) and 1 mM DTT with a Microfluidizer. Proteins were isolated from cleared lysates by GSH affinity chromatography (Serva Electrophoresis), eluted by PreScission protease treatment and further purified by size exclusion chromatography (ENrich™ SEC 70 10/300, Bio-Rad Laboratories, Hercules, CA, United States). Rheb (in 20 mM HEPES pH 7.4, 150 mM NaCl, 2 mM MgCl_2_, 1mM TCEP) was concentrated and stored at -70 °C until use, while the GAP domain (in 20 mM HEPES pH 7.4, 500 mM NaCl, 10 % glycerol, 1 mM TCEP) was used immediately.

### Protein Production in Mammalian Cells

For expression of the different TSC complexes, Expi293F cells (Gibco, Thermo Scientific) were used. Cultures were (co-) transfected with 1 µg DNA/ ml cells and 3 µg PEI/ 1 µg DNA. After 3-5 h at 37°C and 8% CO2, VPA (valproic acid sodium salt, Sigma Aldrich) was added to a final concentration of 3.5 mM. Cells were grown for 3 days before harvest. For purification, 10 ml of TSC purification buffer (40 mM HEPES pH 7.4, 300 mM NaCl, 10 mM MgCl2, 0.3 % CHAPS, 5 % Glycerol, 1 mM DTT and protease inhibitor (HP, SERVA electrophoresis)) per 60 ml culture were added on ice. Cells were resuspended and incubated on a rotating wheel for 15 min at 4°C. The lysates were cleared by centrifugation at 39.000 ×g and 4°C for 30 min, filtered with a 0.45 µm filter and incubated with FLAG M2 affinity resin (Sigma Aldrich). The resin was washed once with the purification buffer containing 10 mM ATP and twice without ATP. FLAG-tagged proteins were eluted with 100 µg/ml FLAG peptide (TargetMol) in purification buffer and elution fractions were supplemented with 1 mM TCEP. For cryo-EM, FLAG peptide was removed by dilution/concentration and buffer was exchanged to TSC cryo buffer (40 mM HEPES pH 7.4, 300 mM NaCl, 10 mM MgCl_2_, 2 % Glycerol; Zeba™ Spin Desalting column, Thermo Scientific).

### GTP Hydrolysis Assay

For the HPLC-based multiple turnover assay, 5 µM GTPase was mixed with 1 mM DTT, 20 mM EDTA and 5 mM MgCl_2_, with or without GAP proteins. To start the reaction, GTP was added to a concentration of 50 µM. At given timepoints, samples were taken, frozen in liquid N_2_ and stored at -70°C. The GTP/GDP ratios were analyzed with an AMAZE HA column (Helix chromatography) on an Agilent HPLC system (Agilent 1260 Infinite II) equipped with an autoloader. Nucleotides were separated on an acetonitrile (Honeywell) gradient and measured at 254 nm. The area under the curve of the GDP and GTP peaks was integrated and the fraction of GTP hydrolyzed was calculated.

### Structure determination

Grids (UltrAuFoil® R 1.2/1.3 on Au 300 mesh grids, Quantifoil Micro Tools GmbH) were glow-discharged (SmartPlasma 2) at 0.1 mbar, 90 s process and 50 % performance, prior to the addition of 4 µl of the protein sample, followed byblotting at 95 % humidity, 14°C, blot force 15 for 5.5 to 6 s and vitrification in liquid ethane. By automated data collection with EPU, 30.379 micrographs without tilting and 17.658 micrographs at 40° tilt angle were collected at a nominal magnification of 230.000× (corresponding to a calibrated pixel site of 0.571 Å) with an accelerating voltage of 300 kV and a total exposure dose of 50 electrons Å^-2^.

Data was processed with CryoSPARC ^26^ (Supplementary Fig. 3). Initial rounds of particle picking were performed with the blob picker followed by picking with crYOLO ^42^. Particles were imported back into CryoSPARC and extracted from micrographs at a box size of 800 pixels and fourier-cropped to a box size of 400 pixels corresponding to a pixel size of 1.142 Å/ pixel to accelerate computations. Multiple rounds of 2D classification were performed, followed by an *ab initio* reconstruction. Promising classes were selected and volumes optimized with homogenous reconstruction, local refinement implemented in CryoSPARC, and 3D classification. A mask for local refinement of the core complex was designed and particles were re extracted at a box size of 800 pixels corresponding to 0.571 Å/ pixel. After extensive processing in CryoSPARC, the particles still suffered from preferred orientation and showed an effective resolution lower than reported by the gold standard FSC_0.143_. Therefore, coulomb potential maps from CryoSPARC sharpening were improved using EMReady2 without providing a model to avoid bias during this process^25^. This substantially reduced preferred orientation and resulted in maps with an effective resolution close to the reported gold standard FSC_0.143_ values.

The TSCC particle shows a pseudo C2-symmetry, which can cause global misalignments especially at the long flexible tails and TSC1. Therefore, a symmetry expansion approach was performed to improve the details of these regions (Supplementary Fig. 6). First, the particles of the two best classes of the ab-initio refinement were pooled and refined with C2-symmetry, followed by C2 symmetry expansion. The symmetry expanded particles were locally refined with C1-symmetry using a mask around half of the TSCC particle, followed by focused 3D classification of the same region. Model building was performed in ChimeraX ^43^ with ISOLDE ^44^ based on AlphaFold2 predictions ^45^ that were fitted into the density and rebuilt.

### mTORC1 activity assay

Functionality of TSC2 variants was tested with a cellular mTORC1 activity assay essentially as described previously ^31^. In brief, HEK293T TSC1/2 dKO cells were transfected with TSC2 (WT or variant), and Myc-tagged TSC1 and S6K constructs using lipofectamine 2000 (Invitrogen). Cells were harvested the next day and cleared lysates analyzed by immunoblotting for levels of TSC1, TSC2, S6K and phospho-S6K (T389). Antibodies used: rabbit polyclonal anti-TSC1 and anti-TSC2 ^31,46^, anti-Myc (Cell Signaling Technology # 2272 and #2276), anti-p70 S6 Kinase-pT389 (Cell Signaling Technology # 9206), IRDye® 680RD Goat Anti-Rabbit IgG and IRDye® 800CW Goat Anti-Mouse IgG (LI-COR).

### Quantification and statistical analysis

For quantification of western blots, data from at least three independent repeats were used. Data are presented as mean ±SD and the significance was calculated using a t-test. Sample sizes (n) and significance values are indicated for each figure panel (* p<0.05, ** p<0.001, *** p<0.0001, **** p<0.00001, n.s. not significant).

## References

1. Wennerberg, K., Rossman, K. L. & Der, C. J. The Ras superfamily at a glance. J. Cell Sci. 118, 843–846 (2005).

2. Vetter, I. R. & Wittinghofer, A. The Guanine Nucleotide-Binding Switch in Three Dimensions. Science (1979). 294, 1299–1304 (2001).

3. Bos, J. L., Rehmann, H. & Wittinghofer, A. GEFs and GAPs: Critical Elements in the Control of Small G Proteins. Cell 129, 865–877 (2007).

4. Inoki, K., Li, Y., Xu, T. & Guan, K.-L. Rheb GTPase is a direct target of TSC2 GAP activity and regulates mTOR signaling. Genes Dev. 17, 1829–34 (2003).

5. Henske, E. P., Józwiak, S., Kingswood, J. C., Sampson, J. R. & Thiele, E. A. Tuberous sclerosis complex. Nat. Rev. Dis. Primers 2, 16035 (2016).

6. Dibble, C. C. et al. TBC1D7 Is a Third Subunit of the TSC1-TSC2 Complex Upstream of mTORC1. Mol. Cell 47, 535–546 (2012).

7. Rasche, R. et al. Structure and mechanism of the RalGAP tumor suppressor complex. Nature Communications 16, (2025).

8. Daumke, O., Weyand, M., Chakrabarti, P. P., Vetter, I. R. & Wittinghofer, A. The GTPase-activating protein Rap1GAP uses a catalytic asparagine. Nature 429, 197–201 (2004).

9. Scrima, A., Thomas, C., Deaconescu, D. & Wittinghofer, A. The Rap-RapGAP complex: GTP hydrolysis without catalytic glutamine and arginine residues. EMBO J 27, 1145–1153 (2008).

10. Fitzian, K. et al. TSC1 binding to lysosomal PIPs is required for TSC complex translocation and mTORC1 regulation. Mol. Cell 81, 2705–2721 (2021).

11. Bayly-Jones, C. et al. Structure of the human TSC:WIPI3 lysosomal recruitment complex. Science Advances 10, 5807 (2024).

12. Yang, H. et al. Structural insights into TSC complex assembly and GAP activity on Rheb. Nat. Commun. 12, 339 (2021).

13. Yoshimura, S. I., Egerer, J., Fuchs, E., Haas, A. K. & Barr, F. a. Functional dissection of Rab GTPases involved in primary cilium formation. Journal of Cell Biology 178, 363–369 (2007).

14. Menon, S. et al. Spatial control of the TSC complex integrates insulin and nutrient regulation of mTORC1 at the lysosome. Cell 156, 771–785 (2014).

15. Capo-Chichi, J. M. et al. Disruption of TBC1D7, a subunit of the TSC1-TSC2 protein complex, in intellectual disability and megalencephaly. J. Med. Genet. 50, 740–744 (2013).

16. Alfaiz, A. A. et al. TBC1D7 mutations are associated with intellectual disability, macrocrania, patellar dislocation, and celiac disease. Hum. Mutat. 35, 447–451 (2014).

17. Schrötter, S. et al. The non-essential TSC complex component TBC1D7 restricts tissue mTORC1 signaling and brain and neuron growth. Cell Rep. 39, 110824 (2022).

18. Ramlaul, K. et al. Architecture of the Tuberous Sclerosis Protein Complex: Tuberous Sclerosis Protein Complex Architecture. J. Mol. Biol. 433, 166743 (2021).

19. Hansmann, P. et al. Structure of the TSC2 GAP Domain: Mechanistic Insight into Catalysis and Pathogenic Mutations. Structure 28, 933–942 (2020).

20. Marshall, C. B. et al. Characterization of the intrinsic and TSC2-GAP-regulated GTPase activity of Rheb by real-time NMR. Sci. Signal. 2, ra3 (2009).

21. Mazhab-Jafari, M. T. et al. An autoinhibited noncanonical mechanism of GTP hydrolysis by Rheb maintains mTORC1 homeostasis. Structure 20, 1528–39 (2012).

22. Fu, W. & Wu, G. Design of negative-regulating proteins of Rheb/mTORC1 with much-reduced sizes of the tuberous sclerosis protein complex. Protein Sci. 32, e4731 (2023).

23. Su, M. Y., Fromm, S. A., Remis, J., Toso, D. B. & Hurley, J. H. Structural basis for the ARF GAP activity and specificity of the C9orf72 complex. Nature Communications 2021 12:1 12, 3786- (2021).

24. Ismail, S. A., Vetter, I. R., Sot, B. & Wittinghofer, A. The structure of an Arf-ArfGAP complex reveals a Ca2+ regulatory mechanism. Cell 141, 812–821 (2010).

25. Cao, H. et al. EMReady2: improvement of cryo-EM and cryo-ET maps by local quality-aware deep learning with Mamba. Nat. Commun. 10.1038/s41467-026-71794-1 (2026) doi:10.1038/s41467-026-71794-1.

26. Punjani, A., Rubinstein, J. L., Fleet, D. J. & Brubaker, M. A. cryoSPARC: algorithms for rapid unsupervised cryo-EM structure determination. Nat. Methods 14, 290–296 (2017).

27. Zech, R., Kiontke, S., Mueller, U., Oeckinghaus, A. & Kümmel, D. Structure of the tuberous sclerosis complex 2 (TSC2) N Terminus provides insight into complex assembly and tuberous sclerosis pathogenesis. Journal of Biological Chemistry 291, 20008–20020 (2016).

28. Lee, W. S. et al. Pathogenic RHEB Somatic Variant in a Child With Tuberous Sclerosis Complex Without Pathogenic Variants in TSC1 or TSC2. Neurology 101, 78–82 (2023).

29. Fokkema, I. F. A. C. et al. The LOVD3 platform: efficient genome-wide sharing of genetic variants. European Journal of Human Genetics 2021 29:12 29, 1796–1803 (2021).

30. Karczewski, K. J. et al. The mutational constraint spectrum quantified from variation in 141,456 humans. Nature 2020 581:7809 581, 434–443 (2020).

31. Dufner Almeida, L. G., et al. Comparison of the functional and structural characteristics of rare *TSC2* variants with clinical and genetic findings. Hum. Mutat. 41, 759–773 (2020).

32. Liu, M. et al. ATR/Chk1 signaling induces autophagy through sumoylated RhoB-mediated lysosomal translocation of TSC2 after DNA damage. Nat. Commun. 9, 4139 (2018).

33. Demetriades, C., Doumpas, N. & Teleman, A. a. Regulation of TORC1 in response to amino acid starvation via lysosomal recruitment of TSC2. Cell 156, 786–799 (2014).

34. Benvenuto, G. et al. The tuberous sclerosis-1 (TSC1) gene product hamartin suppresses cell growth and augments the expression of the TSC2 product tuberin by inhibiting its ubiquitination. Oncogene 2000 19:54 19, 6306–6316 (2000).

35. Santiago Lima, A. J., et al. Identification of regions critical for the integrity of the TSC1-TSC2-TBC1D7 complex. PLoS One 9, e93940 (2014).

36. Gai, Z. et al. Structure of the TBC1D7-TSC1 complex reveals that TBC1D7 stabilizes dimerization of the TSC1 C-terminal coiled coil region. J. Mol. Cell Biol. 8, 411–425 (2016).

37. Qin, J. et al. Structural Basis of the Interaction between Tuberous Sclerosis Complex 1 (TSC1) and Tre2-Bub2-Cdc16 Domain Family Member 7 (TBC1D7). J. Biol. Chem. 291, 8591–601 (2016).

38. Bakula, D. et al. WIPI3 and WIPI4 β-propellers are scaffolds for LKB1-AMPK-TSC signalling circuits in the control of autophagy. Nat. Commun. 8, 15637 (2017).

39. Rasche, R. et al. The GTPase κB-Ras is an essential subunit of the RalGAP tumor suppressor complex. Journal of Biological Chemistry 301, (2025).

40. Winden, K. et al. Tuberous sclerosis complex. Nat. Rev. Dis. Primers 12, 11 (2026).

41. Cheah, P. S. et al. Gene therapy for tuberous sclerosis complex type 2 in a mouse model by delivery of AAV9 encoding a condensed form of tuberin. Sci. Adv. 7, (2021).

42. Wagner, T. et al. SPHIRE-crYOLO is a fast and accurate fully automated particle picker for cryo-EM. Commun. Biol. 2, 218 (2019).

43. Pettersen, E. F. et al. UCSF ChimeraXC: Structure visualization for researchers, educators, and developers. Protein Science 30, 70–82 (2021).

44. Croll, T. I. ISOLDE: A physically realistic environment for model building into low-resolution electron-density maps. Acta Crystallogr. D Struct. Biol. 74, 519–530 (2018).

45. Jumper, J. et al. Highly accurate protein structure prediction with AlphaFold. Nature 596, 583–589 (2021).

46. Hoogeveen-Westerveld, M. et al. Functional assessment of variants in the TSC1 and TSC2 genes identified in individuals with Tuberous Sclerosis Complex. Hum. Mutat. 32, 424–35 (2011).

