## Supplement for "Molecular basis of TSC complex GAP activity"

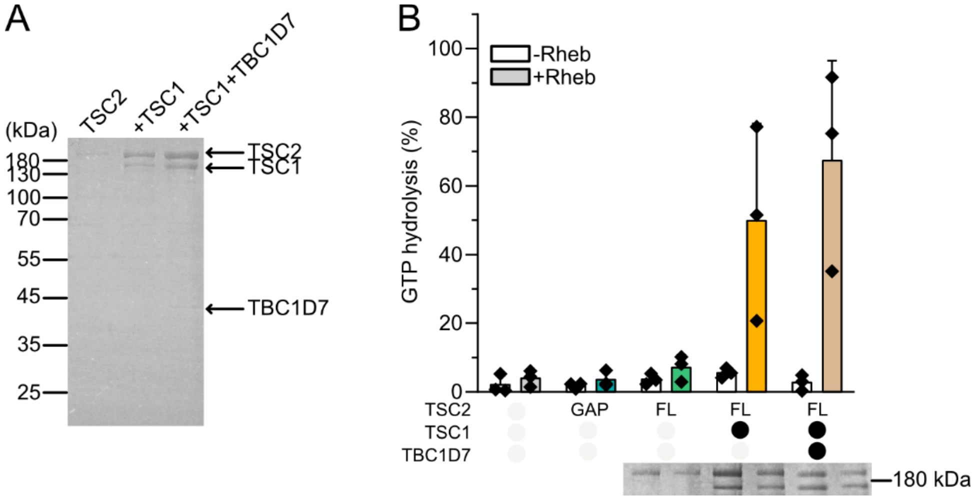


**Supplementary Figure 1: GAP activity of different TSC2 complexes.** (**A**) SDS-PAGE analysis of purified proteins. (**B**) GTP hydrolysis assay data from Figure 1D including controls. Insert shows equivalent TSC2 levels added to the reactions.

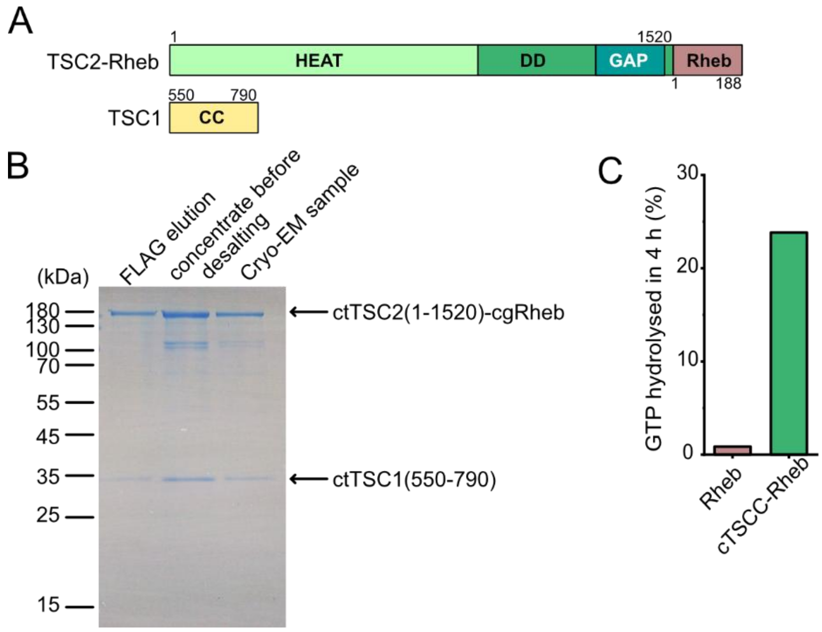


**Supplementary Figure 2: Characterization of a fungal core TSC complex fused with Rheb (cTSCC-Rheb).** (**A**) Architecture of the designed constructs. (**B**) Purification of cTSCC-Rheb for cryo-EM analysis (**C**) GTP hydrolysis assay with cTSCC-Rheb. Equal concentrations of monomeric Rheb and cTSCC-Rheb were assayed.

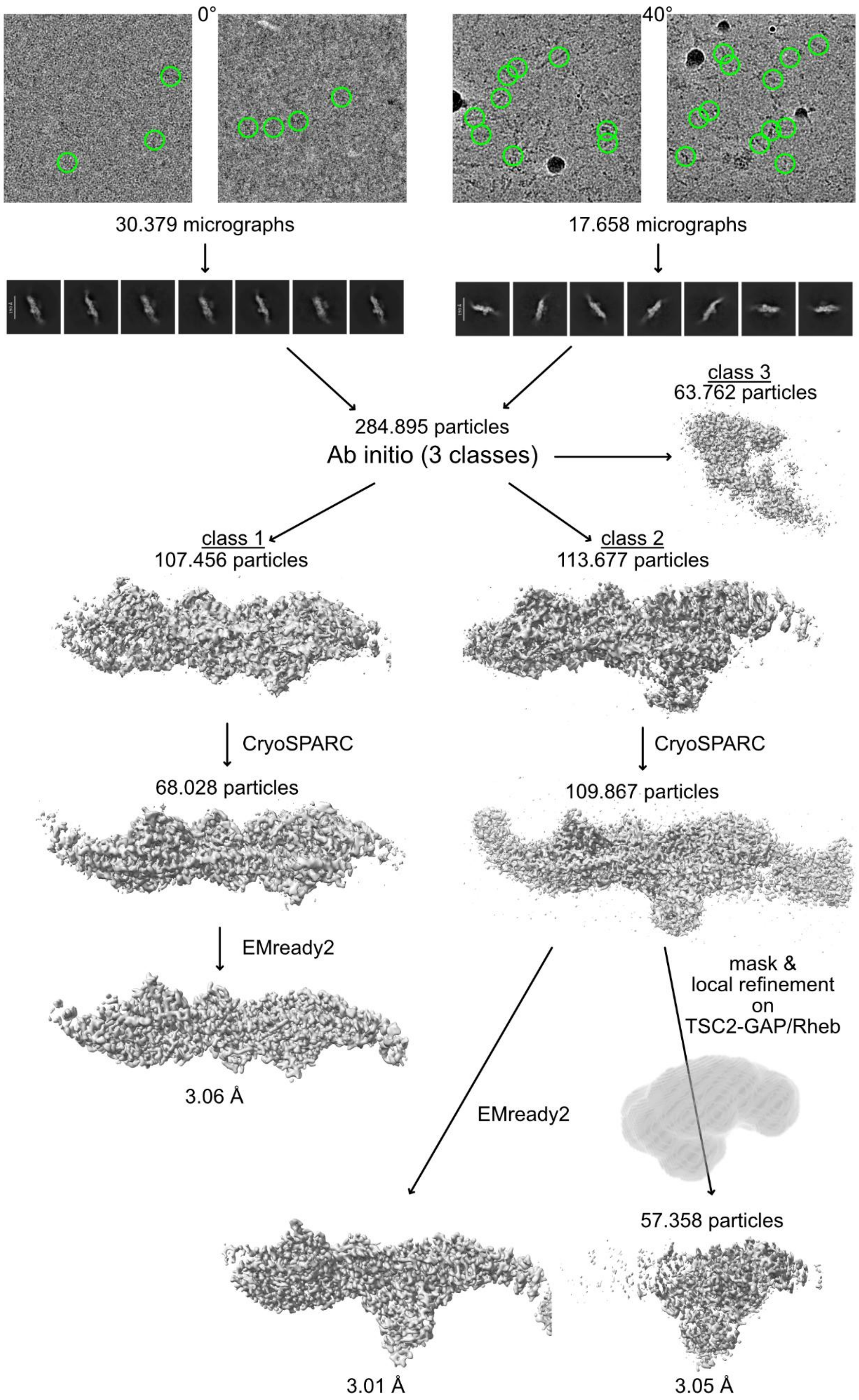


**Supplementary Figure 3: cryo-EM data processing workflow.**

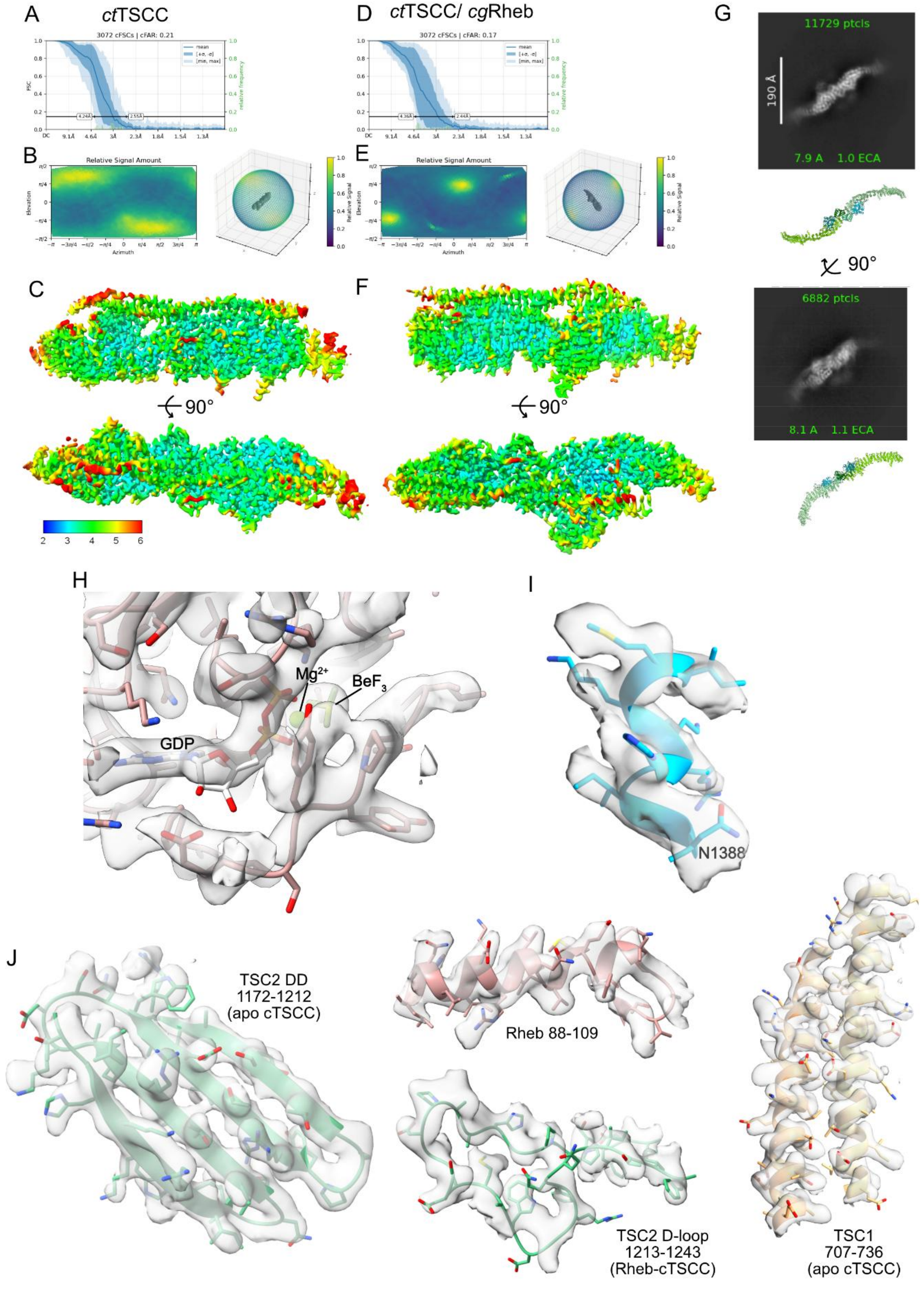
A
ctTSCC
3072 cFSCs | cFAR: 0.21
4.24Å
2.55Å
D
ctTSCC/ cgRheb
3072 cFSCs | cFAR: 0.17
4.36Å
2.64Å
B
Relative Signal Amount
Azimuth
Elevation
E
Relative Signal Amount
Relative Signal
C
F
90°
90°
2
3
4
5
6
G
11729 ptcls
190 Å
7.9 A 1.0 ECA
90°
6882 ptcls
8.1 A 1.1 ECA
H
Mg2+
BeF3
GDP
I
N1388
J
TSC2 DD
1172-1212
(apo cTSCC)
Rheb 88-109
TSC2 D-loop
1213-1243
(Rheb-cTSCC)
TSC1
707-736
(apo cTSCC)

**Supplementary Figure 4: Quality of the cryo-EM reconstructions**. (**A**) FSC of apo cTSCC. (**B**) Orientation diagnostics FSC of apo cTSCC. (**C**) Local resolution distribution of apo cTsCC. (**D**) FSC of Rheb-bound cTSCC. (**E**) Orientation diagnostics FSC of Rheb-bound cTSCC. (**F**) Local resolution distribution of Rheb-bound cTSCC (**G**) Composite model of the cTSCC reconstruction from cryo-EM with the crystal structure of the TSC2 N-terminal domain in comparison to selected 2D classes. (**H**) Close-up of the Rheb nucleotide binding pocket with the experimental map. (**I**) Close-up of the N-TSC2 catalytic helix with the experimental map. (**J**) Selected regions of the cryo-EM maps with modelled structural elements.

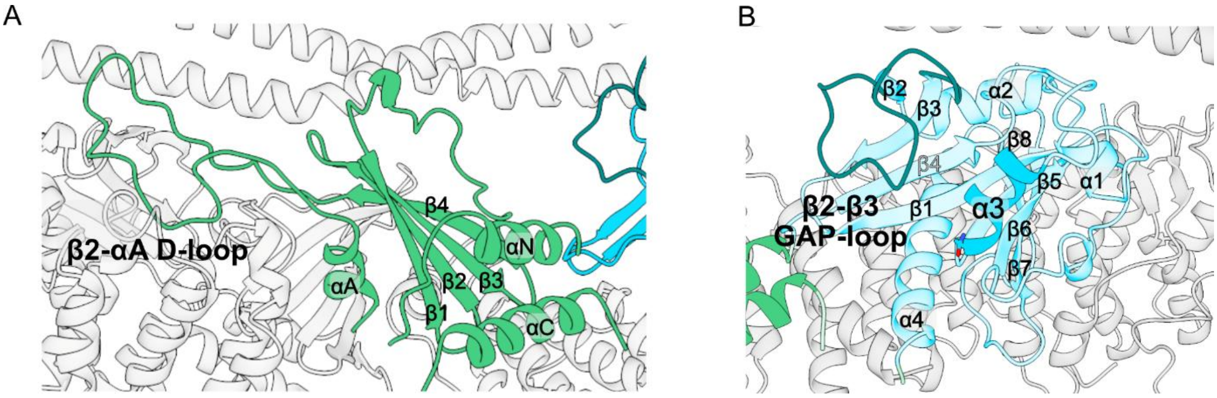


**Supplementary Figure 5: Architecture of apo cTSCC.** (**A**) Organization of the TSC2 dimerization domain. (**B**) Overview of the TSC2 GAP domain fold

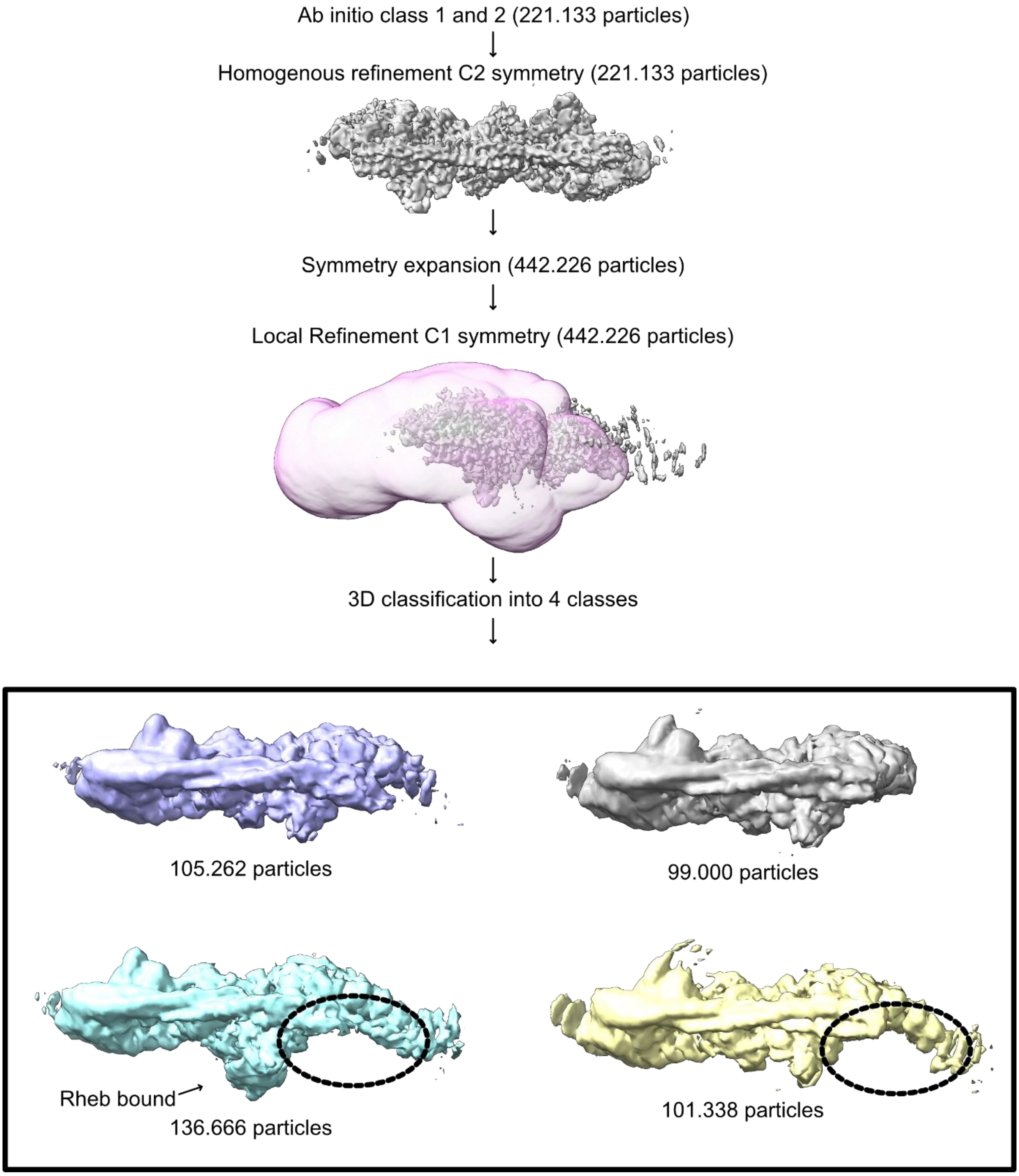


**Supplementary Figure 6: Single particle analysis with symmetry expansion.**

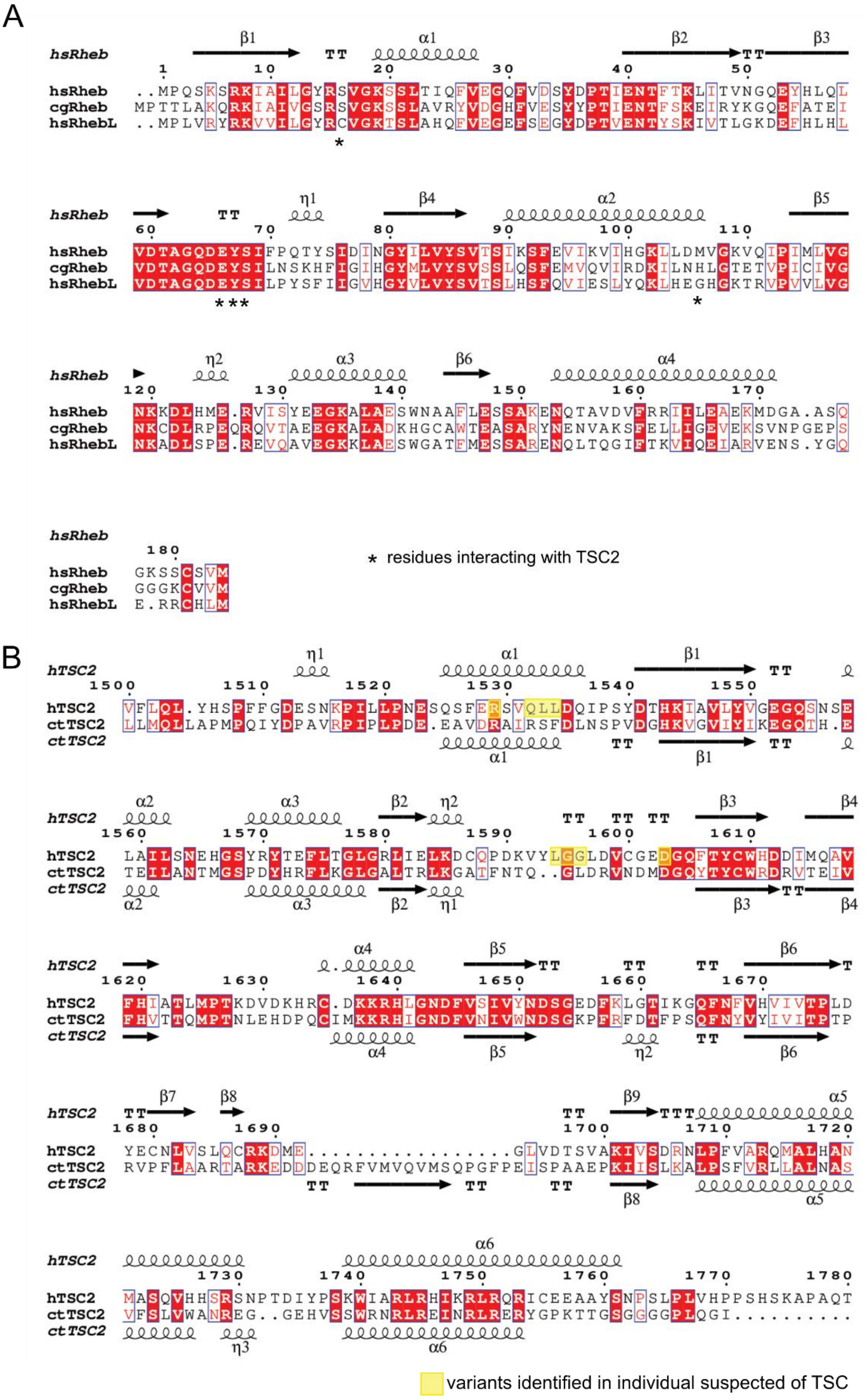


**Supplementary Figure 7**: (**A**) Structure-based sequence alignment of *Chaetomium globosum* Rheb, human Rheb, and human RhebL. Key TSC2 interacting residues are marked by asterisks. (**B**) Structure-based sequence alignment of the *Chaetomium thermophilium* and human αN/αC-helices and GAP domain. Investigated variants are marked by yellow boxes.

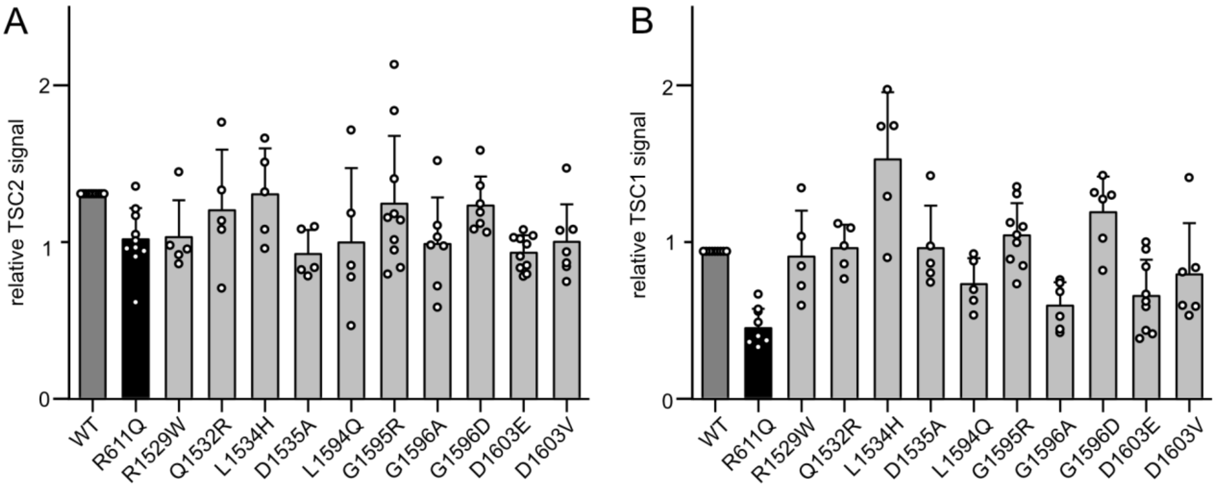


**Supplementary Figure 8**: Quantification of expression levels of (**A**) TSC2 variants and (**B**) TSC1 in reconstitution experiments of TSC1/TSC2 double knock-out HEK293T cells.

**Table S1: Data collection and refinement statistics of cryo-EM structure determination**

| | | |
|---|---|---|
| **Data collection and processing** | | |
| Microscope | Krios G4 with SelectrisX | |
| Detector | Falcon4i detector | |
| Magnification | 210000 | |
| Voltage (kV) | 300 | |
| Electron dose ($e^-$/ $Å^2$) | 50 | |
| Defocus range (µm) | -0,4 to -2,2 | |
| Calibrated pixel size (Å) | 0.571 | |
| Movies/ micrographs | | |
| 0° | 30379 | |
| 40° | 17658 | |
| **Reconstruction** | | |
| Software | CryoSPARC / crYOLO | |
| Total extracted particles (Final Reconstruction) | 68028 | 109867 |
| | *ct*TSCC | *ct*TSCC-*cg*Rheb |
| Number of particles | 68028 | 109867 |
| Symmetry | None | None |
| Final average resolution, gold standard $FSC_{0.143}$ (Å) | 3.06 | 3.01 |
| **Refinement** | | |
| Peptide chains | 4 | 5 |
| Protein residues | 1816 | 1915 |
| Ligands | None | GDP: 1,<br>BeF: 1<br>$Mg^{2+}$: 1 |
| MolProbity Score | 1.2 | 1.3 |
| Clash Score | 0.31 | 0.32 |
| Atoms | 29213 | 30865 |
| rmsd bond distances (Å) | 0.012 | 0.013 |
| rmsd bond angles (°) | 1.956 | 2.067 |
| B-factors (mean) | | |
| Protein | 73.34 | 75.55 |
| Ligand | - | GDP: 51.18<br>$BeF_3$/ $Mg^{2+}$: 69.04 |
| Ramachandran diagram (%) | | |
| Favoured | 96.58 | 95.90 |
| allowed | 3.42 | 4.04 |
| Outlier | 0.00 | 0.05 |
| Rotamer Outlier (%) | 3.00 | 3.32 |

**Table S2: TSC2 variants of uncertain clinical significance investigated in this study.**

| TSC2 residue | structural characterization | variant | source |
|---|---|---|---|
| R1529 | αN helix, facing Rheb D105 | R1529W | LOVD3 (Fokkema *et al.*, 2021) |
| Q1532 | αN helix, facing Rheb D105 | Q1532R | gnomAD (Karczewski *et al.*, 2020) |
| L1534 | αN helix, dimerization domain core | L1534H | LOVD3 (Fokkema *et al.*, 2021) |
| D1535 | αN helix, dimerization domain core | D1535A | LOVD3 (Fokkema *et al.*, 2021) |
| L1594 | β2-β3 loop, Rheb D65/Y67 binding site | L1594Q | LOVD3 (Fokkema *et al.*, 2021) |
| G1595 | β2-β3 loop, Rheb D65/Y67 binding site | G1595R | LOVD3 (Fokkema *et al.*, 2021) |
| G1596 | β2-β3 loop, Rheb D65/Y67 binding site | G1596A | Angela Peron (University of Florence, Florence, Italy) |
| | | G1596D | LOVD3 (Fokkema *et al.*, 2021) |
| D1603 | β2-β3 loop, interaction with catalytic helix α3 | D1603E | LOVD3 (Fokkema *et al.*, 2021) |
| | | D1603V | Luciana Haddad (University of Sao Paulo, Sao Paulo, Brazil) |

**Table S3 Plasmids used in this study**

| Plasmid | Reference |
|---|---|
| pColaHS ctTSC2opt 1285-1491 | (Hansmann *et al.*, 2020) |
| pCDF6P hsTSC2 1538-1729 | (Hansmann *et al.*, 2020) |
| pEF HA hTSC2 isoform 5 | (Zech *et al.*, 2016) |
| pCDF6P GST-hRheb 1-170 | (Hansmann *et al.*, 2020) |
| pCDF6P GST-hRheb 1-170 R15A | (Hansmann *et al.*, 2020) |
| pCDF6P GST-hRheb 1-170 D65A | (Hansmann *et al.*, 2020) |
| pCDF6P GST-hRheb 1-170 Y67A | This study |
| pCDF6P GST-hRheb 1-170 D65A Y67A | This study |
| pCDF6P GST-hRheb 1-170 D105A | This study |
| pcDNA3 3xFLAG hTSC2 | (Manning *et al.*, 2002) |
| pcDNA3 HA *ct*TSC1 550-790 | This study |
| pcDNA3 3xFLAG TSC1 | (Hansmann *et al.*, 2020) |
| pcDNA3 3x FLAG *ct*TSC2 1-1520 | This study |
| pcDNA3 3x FLAG *ct*TSC2 1-1520 *cg*Rheb | This study |
| pcDNA3 HA *ct*TSC1 | This study |
| pOPINEneo-2-Strep-*hs*TBC1D7 | This study |
| pcDNA3 TSC2 | (Hoogeveen-Westerveld *et al.*, 2011) |
| pcDNA3.1 TSC1-Myc | (Hoogeveen-Westerveld *et al.*, 2011) |
| 2B4 Myc-S6K | (Hoogeveen-Westerveld *et al.*, 2011) |
